# Desynapsis in potato is caused by *StMSH4* mutant alleles and leads to either highly uniform unreduced pollen or sterility

**DOI:** 10.1101/2023.02.23.529759

**Authors:** Corentin R. Clot, Dennis Klein, Joey Koopman, Cees Schuit, Christel J.M. Engelen, Ronald C.B. Hutten, Matthijs Brouwer, Richard G.F. Visser, Martina Juranić, Herman J. van Eck

**Author notes:** **Corresponding author** Herman J. van Eck, Plant Breeding, Wageningen University, P.O. Box 386, 6700 AJ Wageningen, The Netherlands.

## Abstract

The balanced segregation of homologous chromosomes during meiosis is essential for fertility and is mediated by crossovers. A strong reduction of crossovers leads to desynapsis, a process in which pairing of homologous chromosomes is abolished before metaphase I. This results in a random segregation of univalent and the production of unbalanced and sterile gametes. However, if desynapsis is combined with another meiotic alteration that restitutes the first meiotic division, then uniform and balanced unreduced gametes, essentially composed of non-recombinant homologs, are produced. This mitosis-like division is of interest to breeders because it transmits most of the parental heterozygosity to the gametes. In potato, desynapsis is a recessive trait that was tentatively mapped to chromosome *8*. In this article, we have fine-mapped the position of the desynapsis locus and identified *StMSH4*, an essential component of the class I crossover pathway, as the most likely candidate gene. A seven base-pair insertion in the second exon of *StMSH4* was found to be associated with desynapsis in our mapping population. We also identified a second allele with a 3820 base-pair insertion and confirmed that both alleles cannot complement each other. Such non-functional alleles appeared to be common in potato cultivars. More than half of the varieties we tested are carriers of mutational load at the *StMSH4* locus. With this new information, breeders can choose to remove desynaptic alleles from their germplasm to improve fertility or to use them to produce highly uniform unreduced gametes in alternative breeding schemes.

## Introduction

Meiosis is a specialized type of cellular division, essential for sexually reproducing organisms, which generates four haploid gametes out of a single diploid mother cell. This is achieved via two successive divisions, meiosis I and II, following a single S phase. While meiosis II resembles a haploid mitosis where sister chromatids are separated, meiosis I is characterized by pairing and subsequent separation of homologous chromosomes. During the prophase of meiosis I, homologous chromosomes pair with each other in a process known as synapsis. The connexion between homologs is maintained until anaphase I by crossovers which ensure their proper positioning and segregation by providing a counterforce to the pole directed spindle forces (Page and Hawley 2003). Indeed, if chiasmata are not formed and synapsis cannot be maintained, such as in asynaptic *Arabidopsis thaliana asy1* mutants, univalents will segregate randomly, ultimately producing aneuploid nonviable gametes (Armstrong et al. 2002). Desynapsis indicates a condition with a similar gametic outcome and is characterized by the failure to maintain synapsis between most homologs after a seemingly initially normal pairing. Desynapsis can result from a drastic reduction in crossover numbers, as observed in organisms with a mutation in the class I crossover pathway (Lynn et al. 2007; Pyatnitskaya et al. 2019). This pathway, also known as ZMM pathway (an acronym for Zip1–4, Msh4–5 and Mer3), is responsible for 75-85% of crossovers in *A. thaliana* (Serrentino and Borde 2012). In addition to their role in chromosome segregation, crossovers are necessary for recombination events which reshuffle parental chromosomes into unique genetic combinations. Although recombination and segregation are essential for genetic diversity, they can pose difficulties for breeders of heterozygous outcrossing crops, as it makes it challenging to preserve previously chosen combinations of beneficial alleles. The highly heterozygous autotetraploid potato (*Solanum tuberosum*) with its slow increase of genetic gains is one embodiment of such a crop. To circumvent this challenge, the potato breeding community is currently putting a lot of efforts in the conversion of the allogamous tetraploid germplasm into a diploid self-compatible germplasm. This new germplasm, compatible with inbreeding, should enable breeding strategies that preserve cumulative genetic gains and delivers true potato seeds (TPS) F1 varieties (Lindhout et al. 2011; Jansky et al. 2016; Zhang et al. 2021; de Vries et al. 2023). Alternatively, if recombination is supressed, as in the case of asynaptic *StDMC1* RNAi mutants (Kumar et al. 2023), the genetic makeup of the parents can be fixed and potentially transmitted to their offspring. This makes it possible to produce TPS without the need for extensive inbreeding. This concept was first explored about three decades ago using desynaptic potato clones that produce unreduced gametes (Jongedijk 1991). Monogenic recessive desynaptic mutants were identified in dihaploids from variety Chippewa and in *S. tuberosum* Group Tuberosum and Phureja hybrids. Those mutations were proven to be allelic and unified under the locus name *Ds-1* (Jongedijk and Ramanna 1988). Locus *Ds-1* was tentatively mapped to chromosome *8* in the bi-parental diploid population CE (Jacobs et al. 1995). Desynaptic mutants are characterised by a 90% reduction in crossover number affecting equally male and female meiosis resulting in the formation of ∼1 bivalent per meiocytes (Jongedijk and Ramanna 1989). Assuming that during meiosis I each univalent has an equal chance of moving to either pole, the probability to obtain a gamete with a normal chromosomal set up is 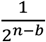 with *n* being the haploid number of chromosomes and *b* being the number of remaining bivalents.

For a desynaptic diploid potato clone, we can estimate that a single reduced gamete out of over 2000 will be balanced, which leads to sterility. However, fertility can be rescued by a second meiotic alteration producing highly uniform unreduced gametes by First Division Restitution (FDR) (Ramanna 1983). Those unreduced gametes, also known as 2n gametes (2nG), are formed when the outcome of meiosis I is restituted by the mis-orientation of meiosis II spindle such as in *ps1* and *jason A. thaliana* mutants (D′Erfurth et al. 2008; De Storme and Geelen 2011). Whether the first division was balanced or chaotic has no impact and meiosis II can progress with the equational division of the entire chromosomal complement. This phenomenon was recently exploited to rescue male fertility in haploid *A. thaliana* (Aboobucker et al. 2023). FDR 2n pollen production is not rare in potato and has been reported in the parents of the population used to map *Ds-1* (Mok and Peloquin 1975; Ramanna 1979).

In the current study, we exploit the joint segregation of desynapsis and FDR 2n pollen to fine map the *Ds-1* locus on the short arm of chromosome *8*. We identified *StMSH4* as candidate gene and discovered a 7 bp insertion in the second exon of the allele associated with desynapsis. Mining the growing number of potato assemblies, we discovered another allele with a 3820 bp insertion at the same position and confirmed that both alleles cannot complement each other. We subsequently found that non-functional *StMSH4* insertion alleles are prevalent in European cultivars. Finally, we discuss the opportunities and limitations offered by desynapsis in the context of potato breeding.

## Materials and Methods

### Plant materials

The diploid mapping population CE-XW comprising 1536 full-sibs descends from a cross between two heterozygous potato clones named C (USW5337.3) and E (77.2102.37) with mixed ancestry of *Solanum tuberosum* Group Tuberosum and Phureja, and *S. vernei*. The seedlings were grown under standard greenhouse conditions at Unifarm (Wageningen University and Research) in 2020 (Clot et al. 2022) and a subset of 500 individuals were grown from tubers the following year. Tubers were planted the 12^th^ of April 2021 in five-litre pots and grown outdoors in a screen cage equipped with sprinkler irrigation. The diploid populations CRH and ERH, used for a complementation test, descend from crosses between the *S. tuberosum* diploid clone RH89-039-16, used as male parent and clones C and E used as female parents. Populations CRH and ERH were sown the 29^th^ of March 2022. For CRH and ERH populations, 100 seedlings were transplanted in 11×11 cm pots and grown in a greenhouse at ambient temperatures and under 16 hours of light.

### Phenotyping pollen stainability and desynapsis

During the flowering stage of the CE-XW population, which occurred between the seventh and the tenth week post-sowing in 2020 and between the fourth and nineth week post-planting in 2021, one pollen sample per flowering individual was collected. Pollen samples were extracted from a freshly opened flower at anthesis using a vibrator pin (modified electric toothbrush). Pollen was spread on a glass slide and stained with a simplified version of Alexander staining (Peterson et al., 2010) and observed under bright field using a Axiophot Zeiss microscope equipped with a Neofluar 10x/0.30 lens. For each sample, four random field views containing approximately 50 pollen grains each were used to visually estimate the proportion of stained pollen relative to the unstained or shrivelled pollen grains. Within the fraction of stained pollen, the proportion of 2n pollen, having a ∼15% larger diameter, was assessed. Combining those two observations, we identified desynaptic individuals. In a wild-type background the effect of desynapsis on pollen stainability is undistinguishable from other forms of sterility. In the presence of another meiotic alteration, resulting in FDR, the unbalanced chromosome segregation of first meiotic division is restored (Fig. 1a). This results in a variable proportion of stained balanced 2n pollen grains next to the shrivelled, unstained, and unbalanced reduced pollen grains. The dichotomous key shown in Fig. 1b allows a binary classification of pollen samples into synaptic and desynaptic classes. Samples with a combination of large stained and small unstained pollen grains were considered desynaptic. Samples with at least 5% of stained pollen grains and including at least one small and stained pollen grain were considered synaptic, irrespective of the level of 2n pollen production.

**Figure 1:**
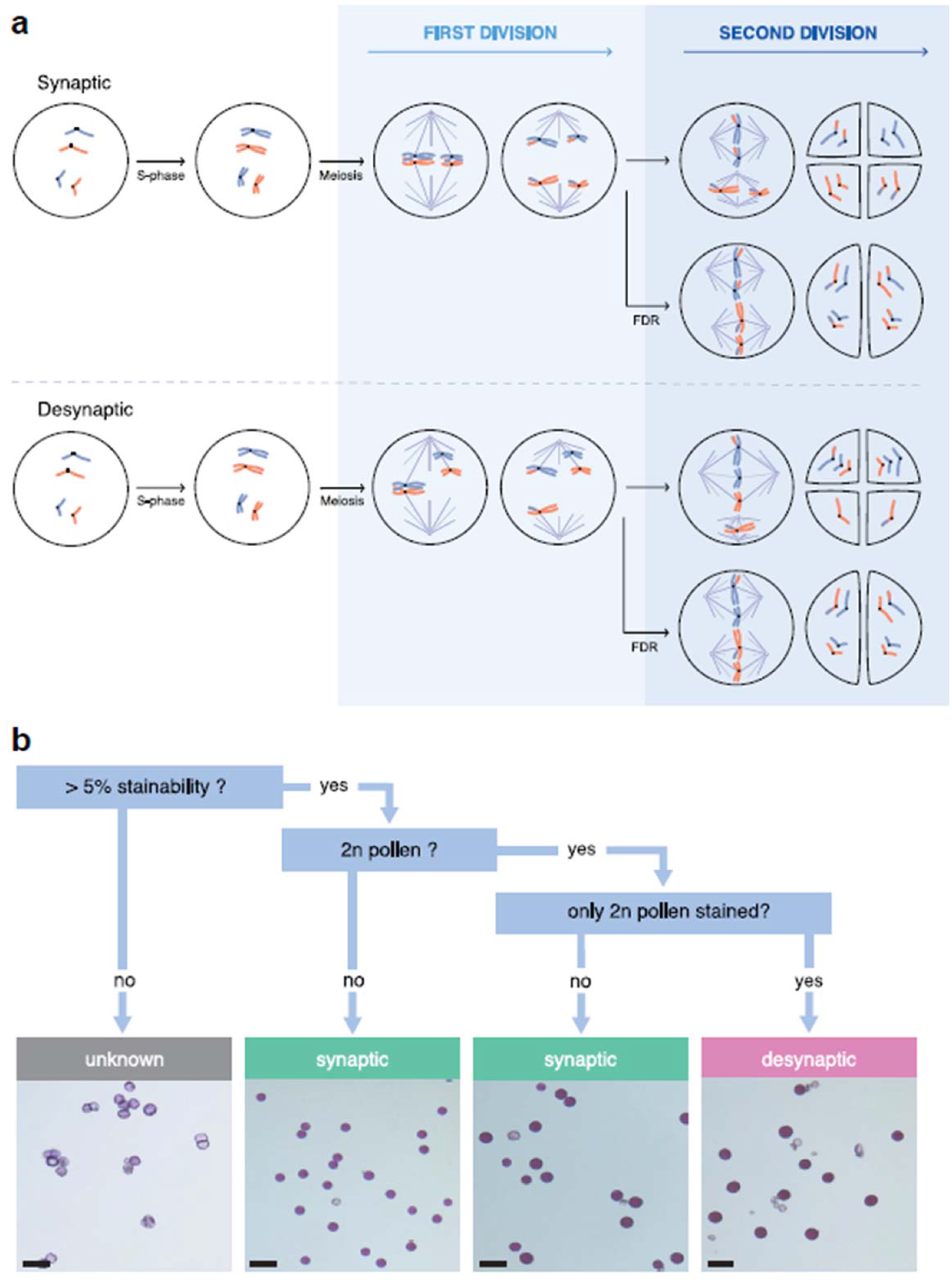
a) Schematic representation of synaptic and desynaptic meiosis producing reduce and FDR 2n pollen. B) Dichotomous key used to phenotype desynapsis in population CE-XW via pollen microscopy. Scale bar is 50 μm.

The same phenotyping protocol, with an increased stringency for synaptic classification set at 5% of stainability of reduced pollen grains, was applied to the CRH and ERH populations which flowered between the nineth and the eleventh week post-sowing. When in doubt about pollen size, pictures were taken with the Zeiss Axiomcam ICc 5 colour camera and pollen diameter was measured with Zen 2.3 lite software. Pollen grains with a diameter above 23 μm were consider as 2n pollen.

### Genetic analysis

Marker data and map construction of population CE-XW is detailed in Clot et al. (2022). A total of 4894 female and 4740 male markers segregating across 1461 individuals were used. QTL mapping was performed using the package polyqtlR version 0.0.6 (Bourke et al. 2018). The function singleMarkerRegression was used to fit an additive model at each marker position returning the -log_10_ p-value of model fit per marker. The significance thresholds for QTL detection were determined via permutation tests on the phenotypic values with N = 1000 cycles and α = 0.05. After an initial QTL discovery using separate maternal and paternal markers, QTL discovery was performed at full classification obtained by merging parental markers of identical physical map position. This full classification was used for the recombinant analysis presented in Suppl. table 1.

**Table 1:**
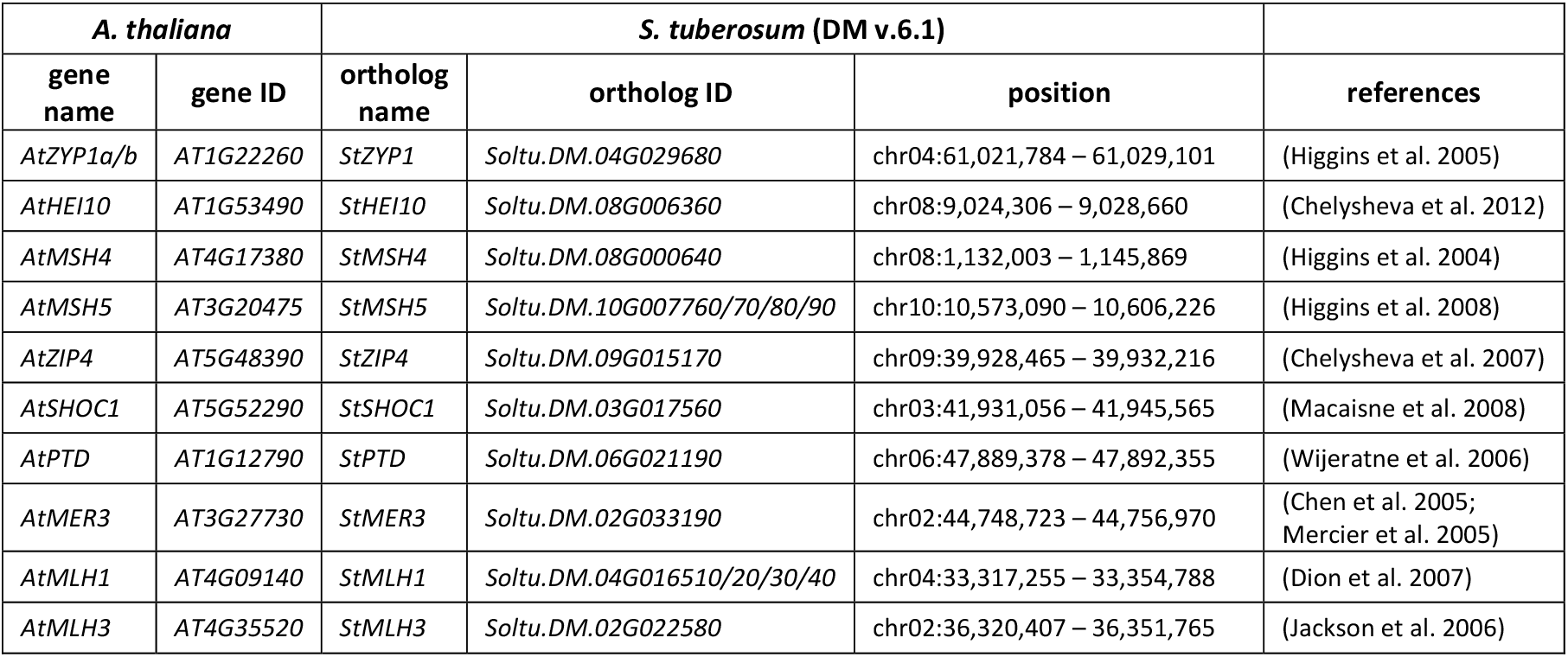
Identification of S. tuberosum orthologs of the A. thaliana ZMM genes including AtMLH1/3 and their positions on the potato reference genome (DM v.6.1)

### Candidate gene exploration

Potato orthologs of *Arabidopsis thaliana* ZMM genes (including *AtMLH1* and *AtMLH3*), necessary for class I crossovers, were identified by blasting the *A. thaliana* protein sequences against DM v.6.1 Protein high confidence gene model on SpudDB (http://spuddb.uga.edu/blast.shtml) with default parameters. Potato orthologs were named according to the *A. thaliana* gene name prefixed by St (for *Solanum tuberosum)* instead of At.

### Retrieving haplotypes of the *StMSH4* locus

Haplotypes of *StMSH4* were mined from a collection of *de novo* assembled potato genomes. A local blast database was build using Nucleotide-Nucleotide BLAST 2.8.1+ (Camacho et al. 2009) with a *S. lycopersicum* cv. Heinz 170 build SL5.0 genome (Zhou et al. 2022) and 18 de-novo assembled potato genomes: M6 v.4.1 (Leisner et al. 2018); DM v.6.1 (Pham et al. 2020); Solyntus v.1.1 (van Lieshout et al. 2020); RH89-039-16 v.3 (Zhou et al. 2020); A6-26, E4-63, E86-69, PG5068, PG6359 and PG0019 (Tang et al. 2022) ; Atlantic v.2.0, Castle Russet v.2.0, Avenger, Atlus, Columba and, Spunta (Hoopes et al. 2022); Otava v.1 (Sun et al. 2022); C88 v.1 (Bao et al. 2022). The genomic sequence of Soltu.DM.08G000640 (*StMSH4* in DM v.6.1) was used as query and the results were manually curated to stitch together partial hits within 5-kb of each other, assumed to be truncated by structura l variations. Incomplete hits matching with scaffold ends were removed from the analysis. Genic and 500 bp upstream and downstream regions from those genome assemblies, homologous Soltu.DM.08G000640, were collected for haplotype analysis. The *StMSH4*.*f1* haplotype, associated with the desynapsis phenotype in our mapping population, was constructed in Geneious Prime 2022.2.2 (https://www.geneious.com) using the merged bam files (ENA BioProject ID PRJEB56778) of the 102 clones phenotyped as desynaptic and genotyped with allelic combination h1h3 at 1.15 Mb on chromosome *8*.

### Multiple sequences comparison

Multiple sequence comparison of *StMSH4* alleles was performed in Geneious Prime 2022.2.2 using MUSCLE 3.8.425 (Edgar 2004) with default parameters. The dendrogram was generated using the neighbor-joining algorithm with default settings.

### KASP marker analysis

Leaf samples of complementation test populations CRH and ERH were sent to Bejo (Warmenhuizen, The Netherlands) for Kompetitive allele-specific PCR (KASP) marker analysis (LGC Genomics GmbH, Berlin, Germany) following to manufacturer protocols. KASP markers distinguishing between putatively functional and non-functional *StMSH4* alleles were developed. Primers are listed in Suppl. table 2. KASP assay results were visualized using SNPviewer (lgcgroup.com/products/genotyping-software/snpviewer) to confirm correct segregation and genotype calling.

### K-mer based exploration of *StMSH4* insertion alleles in commercial germplasm

A *k*-mer based method was used to investigate the presence of *StMSH4* insertion alleles in commercial tetraploid germplasm. Whole genome sequencing reads (150bp pair-ends) of 134 tetraploid potato varieties (unpublished data) were k-merized into 31-mers using KMC v.3.1.0 (Kokot et al. 2017) with default parameters and option -k31. Five additional *k*-mer sets (MSH4_777:T, MSH4_777:C, StMSH4.t1/t2_start, StMSH4.t1/t2_end and, StMSH4.f1 shown in Suppl. table 3) were generated by *k*-merizing sequences of 39 nucleotides overlapping with the T/C polymorphism of KASP marker MSH4_777, sequences of 60 nucleotides centred around both insertion boundaries of the transposon insertion of *StMSH4*.*t1* as well as a sequence of 67 nucleotides centred around the 7 bp footprint of *StMSH4*.*f1*. Those five *k*-mer sets were intersected with the *k*-mer sets of 134 varieties. *K*-mers lacking specificity, present in virtually all varieties, were remove from the analysis which resulted in final sets of 9 *k*-mers specific to *MSH4_777:C*, 9 *k*-mers specific to *MSH4_777:T*, 31 *k*-mers specific to the footprint, 22 *k*-mers specific to the transposon start and, 23 k-mers specific to the transposon end. The absence/presence of *k*-mers in varieties, along with specific k-mers frequencies, were used to infer the distribution of specific *StMSH4* haplotypes in commercial germplasm. The dosage of *MSH4_777:C* was estimated by dividing the *k*-mer frequencies of MSH4_777:C with the sum of MSH4_777:C and MSH4_777:T *k*-mer frequencies.

## Results

### Desynapsis is controlled by a single locus on chromosome *8*

Stained pollen samples from 1345 seedlings of the CE-XW population were observed under a microscope and classified according to the dichotomous key presented in Fig. 1b. These microscope observations allowed classification of 134 individuals as desynaptic and 912 as synaptic while 299 offspring remained unclassified due to either low pollen stainability, low proportion of 2n pollen or uncertainty about pollen ploidy. To validate this classification, a subset of 500 individuals were regrown from tubers the following year, from which 470 pollen samples could be collected. In 107 desynaptic and 333 synaptic clones we observed the same phenotype in both years. Five desynaptic clones were classified as normal in the 2^nd^ year and 25 clones with normal synapsis were classified as desynaptic in the 2^nd^ year. These 30 conflicting classifications (6.3%) were specifically related to individuals with low pollen shed and were discarded from further analysis. Ultimately, we mapped the *Ds-1* locus using 106 desynaptic and 857 synaptic individuals. While the classification of the desynaptic individuals was based on two years data, this was not the case of all synaptic individuals. Nonetheless, we decided to use all genotyped individuals classified as synaptic based on a single year observation, considering that the increase in power offered by a larger cohort will compensate for putative misclassification.

Using this binary phenotypic classification in a single marker regression with female and male markers, we mapped the *Ds-1* locus to the north arm of chromosome *8* at 1.15 Mb both on the female and the male physical map. Merging parental markers of identical physical map positions, resulting in full classification, also localised *Ds-1* at 1.15 Mb. (Fig. 2a-b). A recessive inheritance of desynapsis implies that the recessive phenotype will match only with one of the allele combinations h1h3, h1h4, h2h3 and h2h4. Indeed, three marker classes accurately predict synaptic plants where h1h4, h2h3 and h2h4 are associated with the *ds-1/Ds-1, Ds-1/ds-1* and *Ds-1/Ds-1* synaptic genotypes respectively, with only one or two phenotypic misclassifications, as shown in the mosaic plot (Fig. 2c). The allele combination h1h3 however, indicative of the *ds-1/ds-1* desynaptic genotypes showed 102 true positive and 79 false negative classifications. These false negatives suggest a second 1:1 segregating locus compensating the *ds-1/ds-1* mutants. However, a new QTL analysis for desynapsis within the cohort of individuals with the allelic combination h1h3 at 1.15 Mb on chromosome *8* did not identify such a compensatory locus (Suppl. fig. 1). Upon microscopic re-examination of the slides of 20 random false positive clones, five are better classified as ambiguous due to a low pollen shed, one was confirmed synaptic and 14 were reclassified as desynaptic. Those 14 clones show a stainability above 50% with more than 95% of stained pollen grain being 2n. The occasional observation of a few small and stained pollen grains in those clones were in retrospect also visible in 10 randomly re-examined true positive desynaptic clones. This suggests that desynaptic plants can produce up to 3% of stained small pollen grains, assumed to be haploid or aneuploid cells with a stainable cytoplasm. Overall, false positives were essentially explained by erroneous phenotypic classifications due to a too mild threshold to classify a pollen sample as synaptic based on a single stained small pollen grain, or the potential misidentification of 2n pollen as n pollen in desynaptic clones with elevated levels of 2n pollen production. Finally, after a recombinant analysis in the subset of individuals curated for false positives and with unambiguous recombination breakpoints, we identified the first 1.9 Mb of chromosome *8* as candidate region for desynapsis (Suppl. table 1).

**Figure 2:**
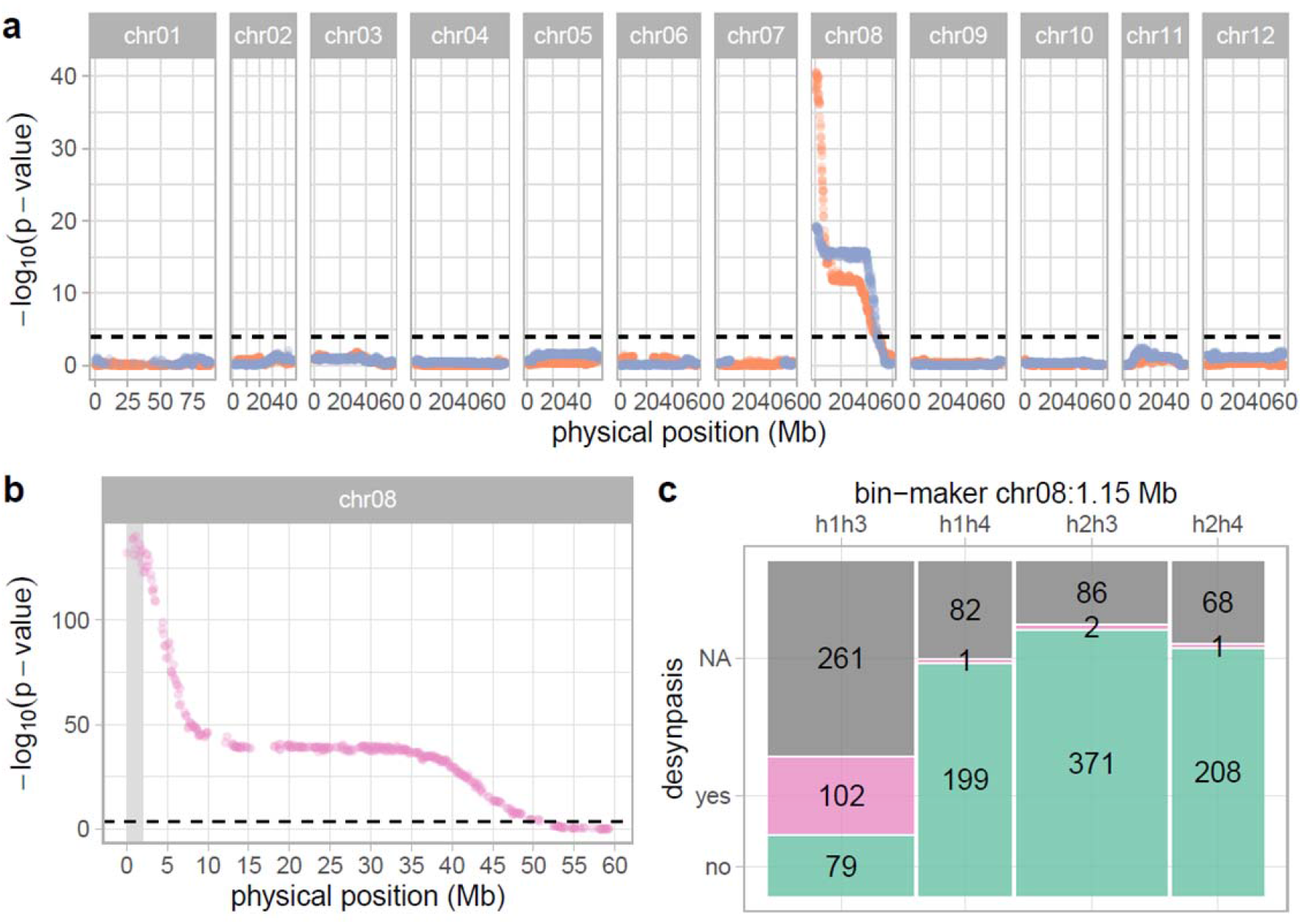
a-b) Significance of the association between markers and the phenotypic classification of desynapsis (n=963). The X axis represents the physical position (Mb), the Y axis represents -log10(p-value) and, the threshold of significance is indicated by the black dashed line. In panel a), clone C marker data are displayed in orange and clone E in blue while in panel b) the integrated marker allele combination h1h3 is displayed in pink and the candidate region for Ds-1 is highlighted in grey c) Mosaic plots illustrating the goodness of fit between phenotypic and genotypic observations for the markers at 1.15Mb on chromosome 8. Individuals phenotyped as synaptic or desynaptic are displayed in green and pink respectively, and phenotypically unclassified individuals in grey.

### Candidate genes in the ZMM pathway

Low amounts of stained n pollen in desynaptic CE material were explained by Jongedijk and Ramanna (1989) with cytological observations. They showed a 90% reduction of crossovers in meiocytes of desynaptic CE plants resulting in unbalanced chromosomal segregation. Class I crossovers, resulting from the ZMM pathway, were shown to represent 75-85% of the crossovers in *A. thaliana* (Serrentino and Borde 2012). Therefore, a gene involved in the ZMM pathway may explain desynapsis in our material. We collected the physical positions of the potato orthologs of *A. thaliana* ZMM genes, including AtMLH1 and AtMLH3, and report their names and positions as found on the potato reference genome DM v.6.1 in Table 1. All *A. thaliana* genes were associated with a single ortholog apart from the duplicated genes *AtZYP1a* and *AtZYP1b* which matched a single gene in DM v.6.1 (*Soltu*.*DM*.*04G029680*) and *AtMSH5* and *AtMLH3* whose orthologs in DM were annotated as four consecutive genes (*Soltu*.*DM*.*10G007760/70/80/90* and *Soltu*.*DM*.*04G016510/20/30/40*). Strikingly, *Soltu*.*DM*.*08G000640* the ortholog of *AtMSH4*, from now on referred to as *StMSH4*, is located around 1.14 Mb on chromosome *8*, in the middle of the candidate region for the *Ds-1* locus. In *A. thaliana, AtMSH4* mutants exhibit a severe reduction in fertility due to an 85% reduction in chiasmata frequency at metaphase I leading to univalents and unbalanced chromosomal segregation (Higgins et al. 2004). This phenotypic description is identical to the phenotype observed in our material.

### Natural diversity of *StMSH4* haplotypes in potato germplasm

Sequence data of *StMSH4*, being a plausible candidate gene for the *Ds-1* locus, were retrieved with BLAST from 18 de novo assembled genomes. This resulted in the identification of at least 19 unique haplotypes. We aligned these sequences with the haplotype of CE-XW desynaptic individuals and calculated a neighbor-joining tree of the haplotypes and rooted the tree using the haplotype from *S. lycopersicum* as outgroup (Fig. 3a). The most common haplotype, *StMSH4*.*1*, was identified in six different clones: DM, RH, Atlantic, PG6359, Otava and, C88. The mutant haplotype associated with desynapsis in our mapping population was identical to the second most common haplotype, named *StMSH4*.*f1* where the f indicates a footprint. This footprint haplotype was found in Colomba, Otava and C88 and is characterized by a seven bp insertion within the second exon of *StMSH4* as annotated in DM v.6.1. Strikingly, much longer insertions of 3820 and 3819 bp are observed at the exact same position in haplotypes named StMSH4.t1 and StMSH4.t2, where the t indicates a transposon insertion (Fig. 3b). These transposon insertion mutants were found in clones RH89-039-16 and Spunta, respectively, and are 99.9% identical with only 10 SNPs and one indel observed within the insertion. Albeit the different insertions observed in haplotypes *StMSH4*.*t1, StMSH4*.*t2* and *StMSH4*.*f1*, the remainder of the haplotypes are identical to each other (Suppl. file 1). We submitted the *StMSH4*.*t1* insert sequence to BLASTn against the RepetDB database (Amselem et al. 2019) “Solanum_tuberosum consensus from ReptDB v2” with defaults parameters. The best hit, with 93% sequence identity, returned the consensus sequence Stub_TedenovoGr-B-G5183-Map16. This sequence is classified as a Class II TIR transposon and is present in 294 copies and 342 fragments in DM v.4.03 and amounts for a cumulative genome coverage of 335,463 bp. In conclusion, we identified *StMSH4* as a candidate gene, and we assume that haplotypes with a premature stop codon due to a transposon insertion or footprint are associated with desynapsis.

**Figure 3:**
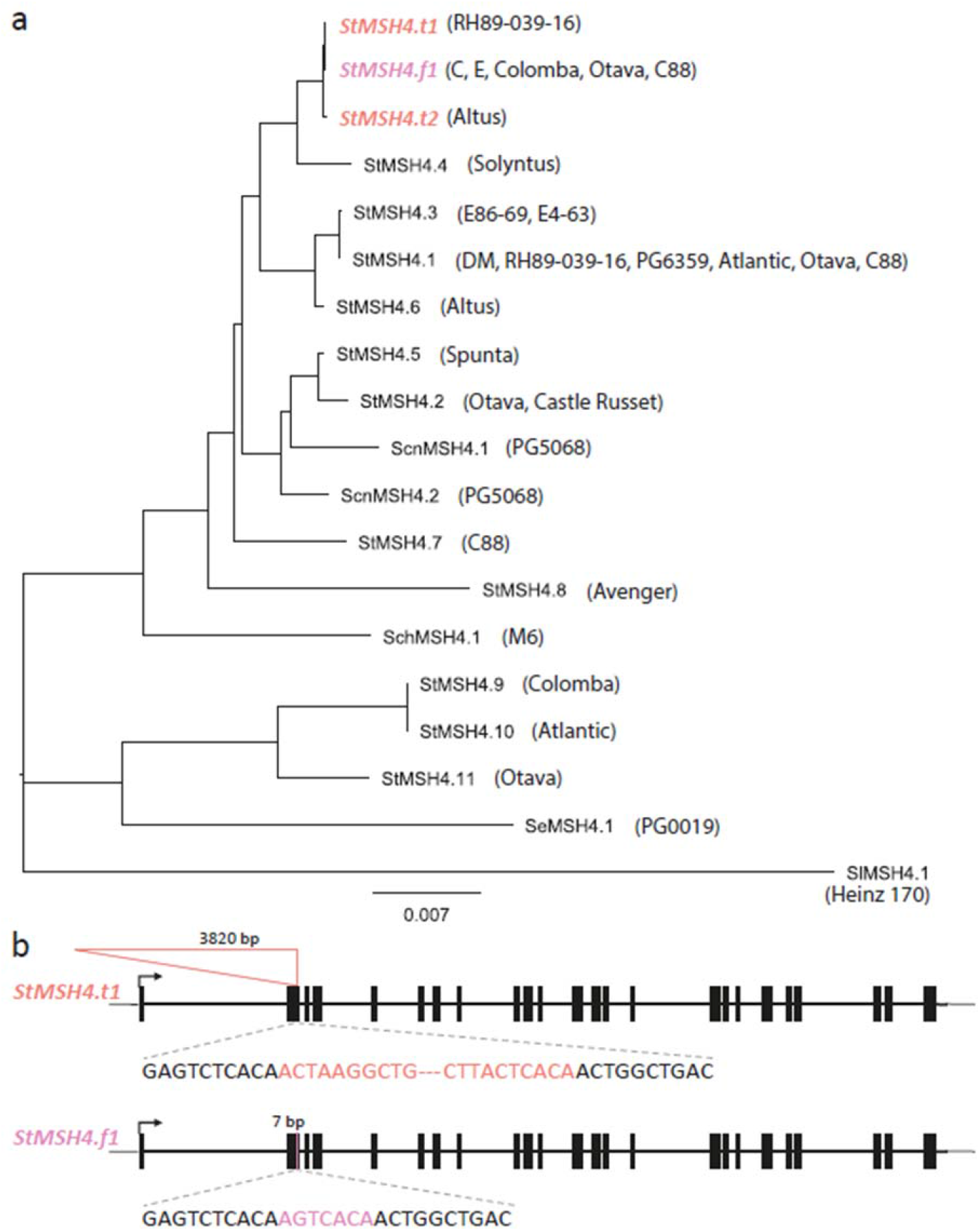
a) Phylogenetic dendrogram of StMSH4 haplotypes retrieved from de novo assemblies. Clone from which the various haplotypes were identified are indicated in between brackets. Haplotypes with insertions in the second exon of StMSH4 are indicates in pink and red. B) Haplotypes StMSH4.t1 and StMSH4.f1 show insertions of 3820 bp and 7bp in the second exon of candidate gene StMSH4.

### Complementation test

The evidence that the *Ds-1* locus is equal to *StMSH4* is only based on a positional co-localisation and phenotypical similarity with *Atmsh4* mutants. To validate our positional evidence with molecular evidence we designed a genetic complementation assay using different insertion mutant versions of *StMSH4* as found in different potato clones. We crossed clones C and E, both heterozygous for *StMSH4*.*f1*, with clone RH89-039-16, heterozygous for *StMSH4*.*t1*, generating populations CRH and ERH. KASP marker MSH4_777 and MSH4_168, both tagging *StMSH4*.*t1* and *StMSH4*.*f1*, were used to follow the segregation of either haplotype in the progenies of CRH and ERH. As expected, in both populations the KASP markers segregated in a 1:2:1 fashion and about one quarter of offspring, homozygous for the KASP alleles associated with the insertion alleles are assumed to carry the haplotype combination *StMSH4*.*t1*/*StMSH4*.*f1*. We phenotyped population CRH and ERH for desynapsis without prior knowledge on the seedling genotypes. Out of CRH 99 seedlings, 47 plants did not flower or shed sufficient pollen for microscopic observation and 3 plants displayed insufficient pollen stainability. Among the remaining 49 plants, 8 were classified as desynaptic and 41 as synaptic. Similarly, for the 100 seedlings of ERH, 49 plants did not flower or shed sufficient pollen for microscopic observation and 3 plants displayed insufficient pollen stainability. Among the remaining 48 plants, 7 plants were classified as desynaptic and 41 plants as synaptic. We observed a perfect correlation between the desynapsis classification and the two KASP markers distinguishing *StMSH4*.*t1* and *StMSH4*.*f1* from the functional haplotypes (Fig. 4). All phenotyped plants bearing the haplotype combination *StMSH4*.*t1*/*StMSH4*.*f1* were classified as desynaptic indicating that *StMSH4*.*t1* and *StMSH4*.*f1* alleles cannot complement each other and are non-functional. From now on, those non-functional alleles can be referred to under the unified notation *Stmsh4*.

**Figure 4:**
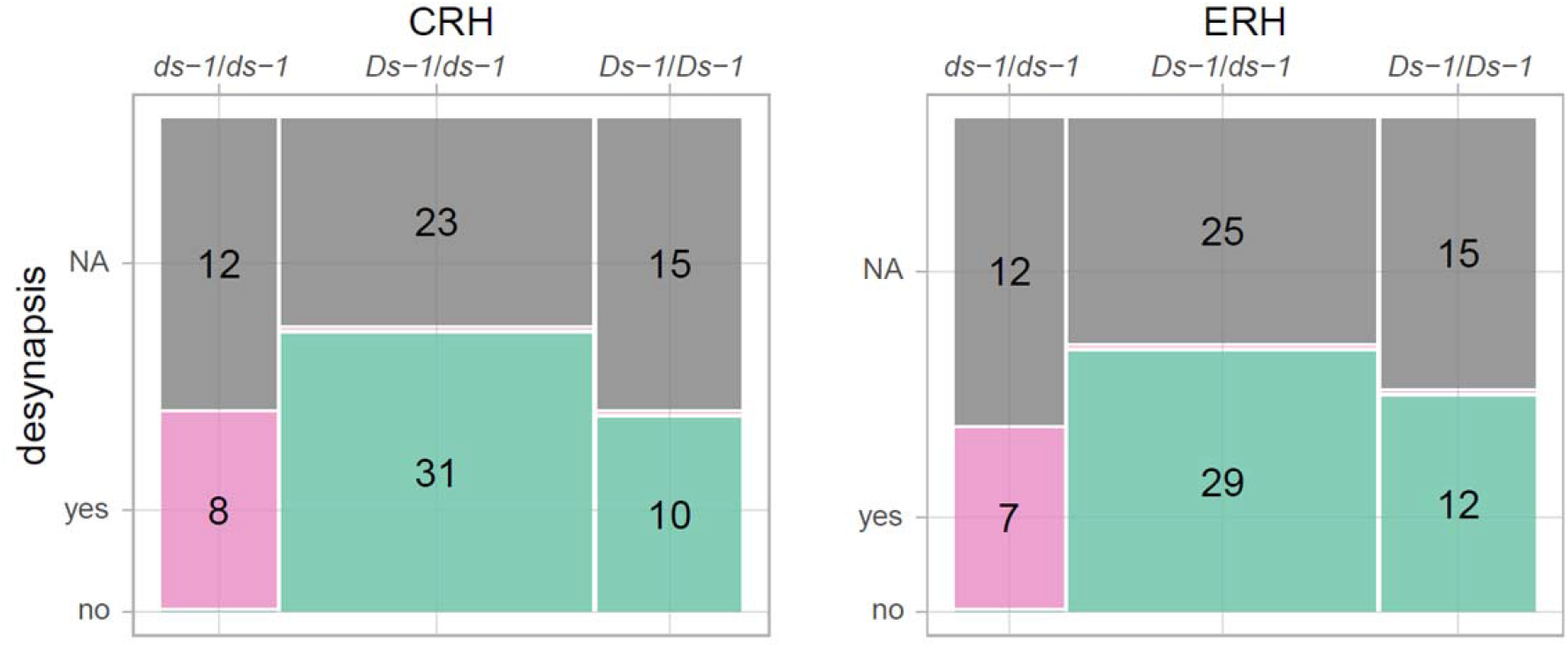
Mosaic plots illustrating the agreement between desynapsis phenotype and genotypes obtained with KASP markers tagging StMSH4 haplotypes in two complementation studies. CRH and ERH indicate the offspring from a cross between clone C or E with clone RH89-039-16. Ds-1 indicates StMSH4 alleles with no insertion and ds-1 stand for either StMSH4.f1 or StMSH4.t1. Individuals phenotyped as synaptic or desynaptic are displayed in green and pink respectively, and phenotypically unclassified individuals in grey.

### Occurrence of *StMSH4* insertion mutants in commercial tetraploid varieties

To assess the occurrence of *Stmsh4* alleles in commercial germplasm, we intersected the *k*-mer sets of 134 re-sequenced tetraploid varieties with the *k*-mer sets uniquely tagging the KASP marker allele *MSH4_777:C*, the footprint region of *StMSH4*.*f1* and the junctions at both ends of *StMSH4*.*t1* transposon insertion site (Suppl. table 3). Surprisingly, we observed at least one of the mutant haplotypes *StMSH4*.*t1* or *StMSH4*.*t2* in 70 out of the 134 varieties, because in these varieties all *k-*mers unique to the junctions of both insertion ends were present. Furthermore, the footprint-specific k-mers were identified in 21 varieties including 14 varieties already positive for the transposon insertion. Two more varieties, Royal and Summer Delight, were found positive for only two k-mers specific for the *StMSH4*.*f1* footprint haplotype. After retrieving reads from these clones containing those two k-mers and aligning them to DMv.6.1, we identified in Royal had a 4 bp footprint instead of the 7 bp insertion found in *StMSH4*.*f1*, indicating a new haplotype called *StMSH4*.*f2* (Suppl. fig 2a). More strikingly Summer Delight retrieved reads mapped partially to the footprint location and partially to another region 28,460 bp downstream. This suggests that the transposon excision event caused a structural rearrangement producing another more complex footprint haplotype (*StMSH4*.*f3*) (Suppl. fig 2b). Whether the *StMSH4*.*f2* and *StMSH4*.*f3* alleles are also non-function remains to be demonstrated. Finally, we observed a perfect correlation between the 79 varieties positive for *k*-mers tagging *MSH4_777:C* and varieties positive for either *StMSH4* insertion mutants. This observation confirms that our KASP marker MSH4_777 accurately predicts the insertion alleles and can be used to estimate their combined dosage. We estimated that a total of 55 varieties were nulliplex for *MSH4_777:C*, while 45 were simplex, 26 were duplex and 8 were triplex. Hence, with a minor allele frequency (MAF) estimated at 22.6%, deleterious *StMSH4* insertion mutants are abundantly present in commercial germplasm.

## Discussion

### Phenotypic classification of desynapsis via pollen microscopy

Rather than relying on laborious cytological observations of male meiocytes to phenotype desynapsis, we exploited the combination of desynapsis and FDR 2n pollen in our population to phenotype this trait through pollen microscopy. Either method has its advantages, where meiocyte observations would allow to capture the reduced chromosome pairing, while pollen observations allow a quick and dirty classification of a large population. A disadvantage of this latter approach is that desynapsis could only be positively classified, and distinguished from other causes of male sterility, when combined with FDR 2n pollen production. This requirement for 2n pollen resulted in 261 unclassified ds*-1*/*ds-1* individuals (59.0%), compared to 236 unclassified *Ds-1*/— individuals (23.2%). Classification of synaptic offspring was highly accurate with only four misclassifications (0.5%), allowing a high mapping accuracy despite 79 misclassifications (43.6%) of desynaptic offspring. We learned that those misclassifications were predominantly caused by two misleading phenotypic observations: 1) almost all stainable pollen grains were unreduced and we lacked small sized pollen grains to see that, and 2) we observed a few stainable small pollen grains while this possibility was not expected in desynaptic clones (initial threshold 0%). Stainability of small pollen grains in a desynaptic individual suggests that balanced segregation may occur by chance alone or that despite an unbalanced genome the cytoplasm is still stainable. This led to a new threshold (<5%), which was successfully applied during the phenotypic classification of the complementation test populations.

### From candidate region to candidate gene

In our mapping experiment, desynapsis was clearly associated with the allelic combination h1h3 on the north arm of chromosome *8*. However, the combined effect of phenotypic misclassification and noisy skim-sequencing data limited our fine mapping resolution. Nonetheless, we performed a recombinant analysis on a curated set of individuals with consistent phenotypic and genotypic data and identified the first 1.9 Mb of chromosome *8* as candidate region for *Ds-1*. Based on DM v.6.1 annotation, this region contains 129 genes among which 7, including *StMSH4*, are putatively involved in reproduction. Articulating the genetic and cytological descriptions of desynapsis in potato (Jongedijk and Ramanna 1989; Jongedijk et al. 1991) with the more recently uncovered function of the ZMM proteins in *A. thaliana* (Pyatnitskaya et al. 2019), we could focus on *StMSH4* as candidate gene for desynapsis. This exemplifies how knowledge obtained from model organism can facilitate discoveries in crops, at least for conserved process such as meiosis.

### Different *StMSH4* insertions alleles

In this study we identified a variety of *StMSH4* alleles, including non-functional alleles due to a transposon insertion in the second exon of this gene or to a 7bp footprint at a position coinciding with the transposon insertion site. Among them, the transposons insertion alleles *StMSH4*.*t1* and *StMSH4*.*t2*, indistinguishable based on our *k*-mer based analysis, were the most widespread (Suppl. table 3). The low sequence divergence between *StMSH4*.*t1* and *StMSH4*.*t2*, limited to 10 SNPs and a 1 bp indel within the insertion sequence, can be explained in three ways. Either those two alleles are *bona fide* and result from 1) independent transposition events at the same location in the same ancestral allele (very unlikely), or 2) from a single transposon insertion event and subsequent divergency restrained to the transposon sequence, or 3) these differences are artifacts caused by the challenge of assembling repetitive sequences present in high copy number in the genome. *StMSH4*.*f1*, the 7 bp footprint allele causing desynapsis in our mapping population, was also relatively common among tetraploid varieties. However, we could also identify two rarer footprint alleles (Suppl. Fig. 2) and cannot exclude the future discovery of more footprints resulting from independent excision events of the widespread transposon alleles.

### *MSH4* and meiotic adaptation to polyploidy

In allotetraploid *Brassica napus*, a reduction in the number of functional copies of MSH4 to a single copy prevents crossovers between homeologous chromosomes without affecting the total number of crossovers (Gonzalo et al. 2019). In this context, reducing *MSH4* dosage seems to help stabilising allopolyploid meiosis by favouring homologous chromosomes as recombination partner. Contrary to allopolyploids, the stabilization of meiosis in autopolyploids, such as *S. tuberosum*, does not rely on recombination partner choice but on avoiding multivalent formation, in particular the combination of trivalent and univalent (Bomblies 2022). This can be mediated by increasing crossover interference strength and ultimately decreasing crossovers number to a minimum of one per pair (Morgan et al. 2021). Interestingly, a direct partner of MSH4, the ZMM protein and synaptonemal complex central element ZIP1 harbour signature of selection for meiotic stabilization in autotetraploid *Arabidopsis arenosa* (Yant et al. 2013). Recent cytological investigation of tetraploid potato cultivars revealed that clone Sante display significantly less multivalent than clones Maris Peers and Cara (Choudhary et al. 2020). Remarkably, Maris Peer and Cara are quadruplex for functional *StMSH4* alleles (Suppl. table 3) and Sante appears to bear one copy of *StMSH4*.*t1* (S. Oome (HZPC), personal communication). Whether reducing the number of functional *MSH4* alleles is an adaptation to auto/allo-polyploidy in potato (i.e to minimize multivalents in autotetraploid *S. tuberosum* or homeologous crossovers in allotetraploid wild relatives such as *S. stoloniferum*) remains speculative but could be simply investigated with functional experiments.

### Desynapsis: a double edge-sword for breeders

At a first glance, the production of sterile n gametes due to desynapsis appears to be a detrimental phenotype that should be removed from potato breeding programs. Suboptimal fertility has always been an issue for potato breeders (Krantz 1924) and maintaining fertility upon inbreeding is central to the reinvention of potato as a diploid inbred line-based crop. With a MAF estimated at 22.6% in tetraploid varieties, *Stmsh4* alleles are contributing to the notorious fertility problems of dihaploids induced from cultivars. Ignoring double reduction, the probability to induce a dihaploid with a given number of *Stmsh4* copies can be calculated using equation (1) with *p* being the ploidy of the parent *P*, P(*P*_n_) being the probability of parent *P* to bear *n* of copies of *Stmsh4* and, P(*G*_k_) being the probability of a gamete *G* (future dihaploid) to bear *k* copies of *Stmsh4*.

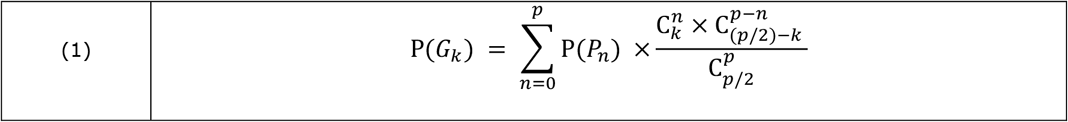

Extrapolating from the estimated dosages of *Stmsh4* in commercial varieties (Suppl. table 3), we can estimate that 6.4% of all induced dihaploids will be homozygous for *Stmsh4* and 33.5% of them will be heterozygous. While these dihaploids are often used to introduce quality and resistance traits to the diploid gene pool, they will also introduce *Stmsh4* alleles, thus hampering future inbreeding efforts. Indeed, one quarter of the S1 population obtained by selfing a clone heterozygous for *Stmsh4* will be desynaptic. Desynapsis will only be observed and selected against when attempting to produce an S2, hereby wasting labour and greenhouse space. We therefore recommend to breeders that aim at developing fertile diploid inbred lines to remove *StMSH4*.*f1* and *StMSH4*.*t1*/*t2* haplotypes from their germplasm.

Looking at desynapsis from another angle, one could envisage to combine it with another meiotic mutation, FDR unreduced gametes (FDR 2nG), and exploit these highly heterozygous and highly uniform unreduced gametes to create tetraploid varieties via sexual polyploidisation. Interploidy breeding schemes exploiting the simplicity of diploid breeding and the heterozygosity offered by tetraploids have long been proposed in potato (Chase 1963; Hutten 1994). In those schemes, tetraploid varieties are produced by unilateral (4x x 2x) or bilateral (2x x 2x) sexual polyploidisation with diploid clones producing 2nG. However, not all 2nG are made equal with FDR 2nG retaining ∼80% of parental heterozygosity compared with ∼40% for 2nG formed by a second division restitution (SDR) (Douches and Quiros 1988a; 1988b; Jongedijk, Hutten, et al. 1991; Peloquin et al. 2008). This higher heterozygosity of FDR 2nG has been linked with a significant yield increase in progenies when compared with SDR 2nG (Kidane-Mariam and Peloquin 1975; Mendiburu and Peloquin 1977; Hutten et al. 1994). In desynaptic clones, the significant reduction in crossover number boosts the heterozygosity of FDR 2nG to 94.1% (Jongedijk, et al. 1991) and thus, could contribute to an even higher heterosis in the tetraploid progeny. Moreover, the increased uniformity of these 2nG will also be instrumental to the development of uniform tetraploid varieties grown from true seeds (TPS varieties), without the necessity to develop inbred parents. Practically, the breeders can generate diploid progenitor material which are heterozygous for *Stmsh4*. These progenitor clones can be used as recurrent parent at the diploid level, but after one generation of selfing the desynaptic descendants producing FDR 2nG can be used for commercial true seed production. Likewise, desynaptic descendants can also be obtained in hybrid progeny descending from a cross between diploid parents that both carry an *Stmsh4* allele.

Despite being less common than FDR 2n pollen production, the formation of FDR 2n megaspores has been reported in desynaptic potato clones by Jongedijk et al (1991). Those authors also discussed the potential use of such mutants to produce heterozygous and uniform tetraploid TPS varieties via bilateral sexual polyploidisation. While being facilitated by the potential of marker assisted selection for desynapsis, further research on the genetic regulation of FDR 2nG production both on the male and the female side remains essential to efficiently exploit desynapsis. Importantly, the genetic regulation of FDR 2nG production is expected to be more complex on the female than on the male side because of the successive type of cytokinesis of female meiosis. FDR 2n pollen could be achieved with a single mutation as in the cases of *Atps1* and *Atjason* mutants (D′Erfurth et al. 2008; De Storme and Geelen 2011). On the other hand, a combination of mutations, such as *Atrec8* and *Atosd1* (d′Erfurth et al. 2009), should be associated with desynapsis to obtain both male and female near non-recombinant unreduced gametes.

### Data availability

The sequencing data of population CE-XW are available from the ENA under the BioProject ID PRJEB56778. The alignment and the raw fasta files of *MSH4* haplotypes, the phenotyping data, genotyping data and, all the R codes necessary to reproduce the results and figures of this article are temporary available at https://figshare.com/s/509dd405f400410a8787, a permanent link with a DOI will be generated for the final version of the manuscript.

### Author contribution statement

CRC: conceived research, collected and analysed phenotypical data, preformed QTL analysis, complementation test, allele mining, wrote manuscript. DK and JK: collected and analysed phenotypical data, performed candidate gene exploration. CS: developed and run KASP assays. CJME and RCBH advised on phenotypical observation, edited breeding discussion. MB provided support for allele mining. RGFV: supervised research, edited manuscript. MJ: supervised research, edited manuscript. HJvE: obtained funding, conceived and supervised research, edited manuscript.

## Acknowledgements

We thank the members of the public-private partnership “A new method for potato breeding: the ‘Fixation-Restitution’ approach” and SusCrop ERANET funded project “DIFFUGAT: Diploid Inbreds For Fixation, and Unreduced Gametes for Tetraploidy” (Averis Seeds B.V., Bejo Zaden B.V., Danespo A/S, Germicopa, Den Hartigh B.V., SaKa Pflanzenzucht GmbH & Co. KG, C. Meijer B.V., and Teagasc) for providing their support. We are grateful to Christian W.B. Bachem, Sara Bergonzi and, Li Shi for their input on *StCDF1* discussion. Our colleagues of Unifarm are acknowledged for plant care in the greenhouse.

